# The mu-opioid receptor gene OPRM1 as a genetic marker for placebo analgesia

**DOI:** 10.1101/139345

**Authors:** Per Aslaksen, June Thorvaldsen Forsberg, Johannes Gjerstad

**Author notes:** Corresponding author: Per M. Aslaksen. Aslaksen and Gjerstad contributed equally to this work. Classification: Biological sciences: Medical sciences; Psychological and cognitive sciences.

## Abstract

The aim of the present study was to examine if genetic factors associated with pain perception could predict the placebo analgesic response in healthy volunteers. 296 participants (182 women) were randomized to either a placebo group receiving placebo cream with information that the cream was an effective painkiller, or to a natural history group receiving no treatment. Pain was induced by contact heat stimuli. Genotyping for the mu-opioid receptor gene OPRM1, the serotonin transporter gene 5-HTT, and the dopamine-metabolizing gene COMT was performed. Individuals with the OPRM1 A/A genotype reported significantly higher placebo responses compared to individuals with the */G variant. No clear effect of the 5-HTT or COMT was observed. The OPRM1 A/A had a predictive accuracy of 92.5% in identification of placebo responders. Our data indicate that the OPRM1 rs1799971 A/A genotype can be used as a reliable identification marker for placebo analgesia.

## Introduction

Several biological and psychological factors may influence the magnitude of placebo responses, and whether individuals experience placebo responses or not (1, 2). Knowledge of which subjects are likely to display the placebo response also provides the benefit of less statistical noise in drug trials and enhance the power of randomized trials (3). Exclusion of placebo responders from drug-trials might reduce the costs of drug development and testing by allowing for smaller sample sizes (4).

The way in which a placebo affects the experience of pain has been extensively studied in both experimental and clinical settings (5–8). However, previous attempts to a priori identify placebo responders by using individual factors as predictors for placebo analgesia have so far had limited success. General personality factors and psychological traits connected to both expectations of drug effectiveness and the emotional aspects of pain relief do not necessarily capture the variability in states that influence whether a placebo is effective or not. Trait measures that are stable even in different contexts, such as genetic composition, are probably more suitable to be predictors for placebo analgesia (4). Hence, identification of genetic markers for placebo responders may be important for the design of future clinical studies, and in clinical settings, recognition of placebo responders may be important for how health-personnel communicate with patients (9) and for the dosage of therapeutic agents (4).

The placebo analgesic response is shown to be modulated by cognitive and emotional processes towards pain and the anticipation of pain (10, 11). Thus, in addition to the sensory aspect of pain experience, predictors of the placebo analgesic response should also be predictive for the affective component of pain sensation measured concomitantly with pain intensity levels.

Previous studies have shown that opioid antagonists may reverse placebo analgesia (12–14). Therefore, placebo analgesia is likely linked to activation of the endogenous opioid system. Moreover, genetic variability related to the function of this system could also affect the placebo response. One such genetic factor is the single nucleotide polymorphism (SNP) called All8G rsl799971 in the opioid receptor mu 1 (OPRM1) gene. This SNP leads to a substitution of asparagine (Asn) to aspartic acid (Asp) at amino acid 40 and subsequent removal of a putative N-linked glycosylation site in the receptor (15). Earlier findings suggest that this SNP leads to reduced OPRM1 N-glycosylation and decreased stability of the receptor in cell culture (16). Moreover, genetic variability that influences the function of the 5-HT transporter (5-HTT) that controls 5-HT re-uptake might be of importance for experimental pain (17). Several genetic variants in the 5-HTT gene SLC6A4 affect 5-HTT expression. One of these is the 5-HTT promoter repeated-length polymorphic region (LPR) (18). Two common allelic variants have been described, a short (S) allele of 14 repeats and a long (L) allele of 16 repeats (19). The S allele seems to reduce the transcription rate and lower expression of the transporter in cell membranes (20, 21). There is also a SNP in the 5-HTT SLC6A4 gene, an A to G substitution rs25531, in the promoter (18, 22). Both the LPR and SNP rs25531 may affect the function of the 5-HTT.

In addition, several lines of evidence show that the SNP in the gene encoding catechol-0- methyltransferase (COMT), Val158Met rs4680 affects sensory processing (23, 24). Individuals homozygous for the Met-allele have a 3-4 times reduced enzyme activity compared to those homozygous for the Val-allele (25). Thus, SNP rs4680 may be associated with sensitivity to experimental pain (23, 26). Moreover, evidence exists that genetic variability in the gene encoding COMT may be important for the development of hyperalgesia and pain catastrophizing (27). Hence, the COMT Val158Met rs4680 also might affect pain modulation and placebo analgesia.

Taken together, genetic variability in genes encoding OPRM1, 5-HTT and COMT may be crucial to understand the neuronal processing important for hyperalgesia (28) and affective responses to pain and pain anticipation (27, 29). Thus, we hypothesized that the genetic variants mentioned above were significantly predictive for the placebo analgesic response. Specifically, we expected that the OPRM1 should be predictive for the placebo analgesic response, and that the 5-HTT and COMT genotypes should be related to the affective components of placebo responding.

The present study was designed as an experimental study with repeated measurements consisting of a calibration of the pain intensity followed by two pretests, a treatment period, and three posttests. Thus, pain was measured in five trials. Experimental pain reports shows substantial inter-individual variability (30, 31), and the calibration procedure was performed to produce equal pain levels in the pretests. The participants were randomized to either a placebo group where a placebo cream was applied together with information that the cream was an effective painkiller, or to a natural history group performing the same pain stimulations as the placebo group, but with no application of cream or information about expectations of pain experience. To ensure blinding of the experimenters, a third group (n=31) where participants received a true anesthetic cream (lidocaine/prilocaine) was included, but data from this group were not used in the analyses. Pain intensity was reported during each pain stimulation by a computerized visual analog scale. Affective responses and blood pressure were measured before the pretests, and after the pretests, after the treatment period, and after the posttests. Saliva samples for genotyping were obtained after the last posttest.

## Results

Table 1 shows the frequency of carriers of the genotypes included in this study. A linear mixed model (LMM) with pain intensity as the dependent variable and the repeated factor Trial was performed. Specifically, the main effects of Trial, Group, OPRM1, COMT, 5-HTT, and sex ofthe participant were included. The interactions were Trial by Group, Trial by OPRM1, Trial by COMT, Trial by 5-HTT, Group by OPRM1, Group by COMT, Group by 5- HTT, Group by Sex, and the three-way interactions between the genetic variables (OPRM1, COMT, 5-HTT), Group and Trial. Figure 1 displays the repeated data. The OPRM1 was dichotomized into OPRM1 *A/A* versus OPRM1 A/G+G/G (*/G) (28). For the data regarding 5-HTT and COMT, however, a three allelic model was used in the analyses (5-HTT *S/S* versus 5-HTT S/L_G_+L_A_/L_A_+S/L_A_ versus 5-HTT L_A_/L_A_ (17) and COMT Val/Val versus COMT Val/Met versus COMT Met/Met) (24). Data is available at http://datadryad.org/review?doi=doi:10.5061/dryad.dd421 (32).

**Figure 1.**
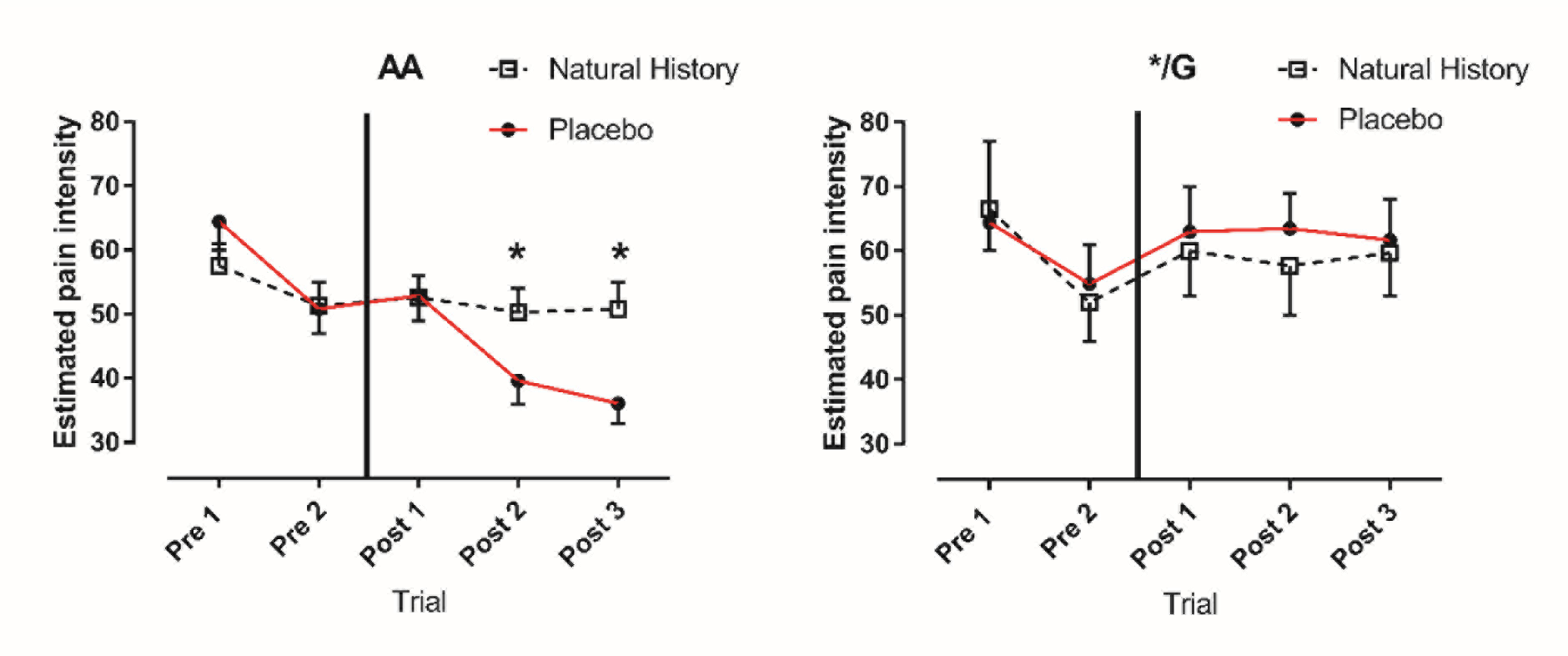
Repeated measures data for Group X OPRM1 (A/A, */G) by Trial. Estimated pain intensity on a 0-100 visual analog scale based on the linear mixed model. Error bars represent 95% confidence intervals. * represent significant Bonferroni-adjusted differences (both p < .01) between the placebo and the natural history group. The x-axis shows the trials and the vertical line specify when the placebo manipulation was performed.

**Table 1.** Frequency of OPRM1, COMT and 5-HTT alleles in the sample of296 participants allocated to the placebo group and the natural history group.

|  | OPRM1 |  | COMT |  |  | 5-HTT |  |  |
| --- | --- | --- | --- | --- | --- | --- | --- | --- |
|  | A/A | */G | A/A | A/G | G/G | S/S | S/L G, L/L<br>A/G, S/L A | L/L, A/A |
| Frequency/participants | 223 | 65 | 92 | 138 | 60 | 59 | 123 | 62 |
| Total number of<br>samples successfully<br>genotyped (%) | 288 (97) |  | 290 (97) |  |  | 244 (82) |  |  |

### Pain

There was a significant main effect of OPRM1 on pain intensity (F (1, 220.67) = 12.49, p <.001) with A/A participants reporting lower pain than G/* participants. Neither 5-HTT (F (2,221.41) = .37, p = .69) nor COMT (F (2, 220.86) = .28, p = .75) had significant main effects on pain in this design. Pain reports changed significantly across Trials (F (4, 556.85) = 33.05, p <.001), see Figure 1.

The placebo effect was significantly stronger in participants with the A/A variant of the OPRM1 gene, shown by the Group by Trial by OPRM1 interaction (F (4, 566.89) = 9.04, p < .001). Bonferroni-corrected post hoc tests showed that the difference between A/A and */G subjects was significant (p < .03) during the last two trials after the placebo manipulation. Males reported lower pain than females (F (1, 220.971) 1.49, p.001), but there were no differences between males and females regarding the placebo effect shown by the interaction of Group by Sex (F 1, 220.97.)83, p.36). As assumed, the participants exhibited significant individual variance, shown by the significant covariance parameter of the intercept of subjects (Variance = 293.54 [95% CI: 238.02-360.72], Z = 9.49, p < .001). A second LLM was fitted by removing the non-significant variables COMT and 5-HTT. This model yielded similar results as the first, but the fit to the data was poorer, as indicated by a worse model-fit (Akaikes Information Criterium; AIC 10491) compared to that in the first model (AIC = 9279). The same significant effects and interactions were present in the reduced model and the first model that included COMT and 5-HTT. The baseline measures of pain, stress and systolic blood pressure were not significantly different between the A/A and */G participants, see Table 2.

**Table 2.** Overview of the sample by OPRM1. *)Calibrated temperature Temperature needed to evoke 60 points on the 0-100 visual-analog scale for pain intensity. BPBlood pressure.

|  | OPRM1 A/A |  | OPRM1 */G |  |  |
| --- | --- | --- | --- | --- | --- |
|  | Mean | 95% CI | Mean | 95% CI | P |
| Age | 21.9 | 21.6-22.4 | 21.9 | 21.3-22.6 | .99 |
| Calibrated temperature for VAS<br>= 60* | 47.5°C | 46.9-48.1 | 47°C | 46.2-47.8 | .37 |
| Stress before procedure | 3.7 | 3.5-3.9 | 4.1 | 3.6-4.6 | .17 |
| Systolic BP before procedure | 130.1 | 128-132.1 | 133.2 | 128.7-137.6 | .16 |

### Subjective stress and blood pressure

Two separate LMMs were performed on the change in stress and SBT (post-pre). Thus, these analyses were performed as univariate analyses without the repeated effect. In the stress data, there was a significant main effect of COMT (F (2, 1101)4.05, p.018) on the change in stress, with COMT AA carriers displaying a higher reduction in self-reported stress than COMT AG carriers (p = .01 Bonferroni adjusted), whereas there was no significant difference between AA versus GG or GG versus AG. Males reported a higher stress reduction than females (F (1,11012) 2.61, p < .001). There was a main effect of 5-HTT on the stress change (F (2, 1101) 4.70, p.009), with SS carriers reporting higher stress reduction (p = .018, Bonferroni adjusted) than S/L_G_+L_A_/L_G_+S/L_A_ carriers, while there were no differences between SS versus L_A_/L_A_ or L_A_/L_A_ versus S/L_A_+L_A_/L_A_+S/L_A_. There was a main effect of OPRM1 on stress change (F (1, 11013) .95, p.047), and the Group by OPRM1 interaction was significant (F (1, 1101) 5.65, p.018) with A/A carriers in the placebo group reporting higher stress reduction than G/* carriers (p = .038, Bonferroni adjusted).

In the SBT data, there were no main effects of 5-HTT, COMT or OPRM1 on the change in SBT. However, the Group by 5-HTT interaction (F (2, 11011) 5.54, p < .001) showed that L_A_/L_A_ carriers in the placebo group showed a higher decrease in SBT than S/L_A_+L_A_/L_A_+S/L_A_ carriers (p = .01 Bonferroni adjusted), whereas there was no other difference in the placebo group. In the natural history group, L_A_/L_A_ carriers displayed a higher reduction in SBT than S/L_A_+L_A_/L_A_+S/L_A_ individuals and S/S carriers (both p’s < .001 Bonferroni-adjusted). The Group by COMT interaction (F (2, 11014) .44, p.012) was significant with subjects in the natural history group displaying a higher SBT decrease if they were *AI*A carriers than both A/G (p = .024) and *GIG* (p.004) carriers shown by Bonferroni-corrected post hoc tests. Thus, the effect of COMT was merely on SBT reduction, not placebo responding per se.

### Prediction of placebo responding

To assess the predictive value of OPRM1 on the placebo analgesic response, we performed a logistic regression analysis on data from the Placebo group (Table 3). The dependent dichotomous variable was stated as whether a reliable placebo effect had occurred from the first pretest to the last posttest, defined by a change in pain scores of more than 13 points on the 100- point VAS scale, which has been used to define a clinically significant change in pain (33, 34). Of the 162 participants in the placebo group, 98 (27 males) displayed a decrease in pain of 13 VAS points or more. The classification accuracy of OPRM1 to identify placebo responders was 92.5%, and the ability to classify non-responders was 48%, giving a total accuracy of 76.9% (Table 4).

**Table 3.** Logistic regression analysis with OPRM 1 SNP as predictor for placebo responding. Placebo responding was defined as a change in pain of more than 13 points on the VAS scale after placebo administration. Explained variance was 27% indicated by Nagelkerke R-square.

|  | B | S.E | Wald | df | P | OR | 95% CI<br>for OR |
| --- | --- | --- | --- | --- | --- | --- | --- |
| OPRM1 (A/A = 1, */G = 0) | 2.43 | .48 | 25.14 | 1 | < .001 | 11.34 | 4.4-29.3 |
| Constant | -1.23 | .43 | 8.23 | 1 | .004 | .29 |  |

**Table 4.** Classification table for the logistic regression analysis for the placebo group. The presence of the A/A variant of the OPRM1 SNP was coded as “1”, whereas presence of the */G variant was coded as “0”.

|  | Predicted |  |  |
| --- | --- | --- | --- |
| Observed | Placebo non-responder | Placebo responder | Percentage correct |
| Placebo non-responder | 24 | 26 | 48 |
| Placebo responder | 7 | 86 | 92.5 |
| Overall percentage |  |  | 76.9 |

## Discussion

The present study showed that individuals with OPRM1 A/A reported significantly higher placebo responses than individuals with OPRM1 */G. Hence, the OPRM1 genotype was associated with the placebo analgesic response. In contrast, no clear effect of the 5-HTT or COMT genotype on placebo responding was observed in the present study.

Interestingly, the logistic regression model performed on those who received the placebo treatment suggested that OPRM1 A/A carriers had an approximately 11 times higher probability of reporting a placebo response than OPRM1 */G carriers. Thus, our results suggest that identification of placebo responders, when heat is used as a test stimulus, may be predicted by OPRM1 genotyping. On the other hand, the lab procedure used in the present study and subsequent analyses of the data was less effective at identifying non-responders. Furthermore, the explained variance in the logistic regression was 27%, leaving a substantial proportion of unexplained variance for the selected model.

Our study support the earlier observation that placebo analgesia is partly dependent on activation ofthe endogenous opioid system (13, 35). Moreover, in line with Pecina et al. (36), we found that OPRM1 A/A carriers had higher placebo responses than OPRM1 */G carriers. Hence, the role of the opioid system in placebo analgesia seems to be replicable. However, the reduced placebo response in the */G carriers could also be related to reduced placebo-related dopaminergic neural transmission or an interaction with other signaling related to 5-HTT or COMT (37,38).

Experimental pain reporting varies across healthy individuals (30, 31). A possible way to reduce this variability is calibration of the stimulus intensity before the experimental procedure (39), as performed in the present study. Nonetheless, pain ratings in both groups still showed substantial intra-individual variability across trials. Moreover, the experience of pain may also depend on the modality of the test stimulus. Although only the OPRM1 genotype seemed to predict placebo analgesia following heat stimuli, the possibility that the 5-HTT or COMT genotype may be predictors following other pain stimuli cannot be excluded.

All participants in our study were healthy volunteers with no history of chronic pain and might, therefore, have different expectancies of a drug effect regarding analgesics compared to pain patients in need of pain relief. In addition, patients enrolled in an RCT probably display higher emotional engagement regarding hope for improvement and the desire for relief (40) compared to participants in an experimental pain study. Thus, as shown in several experimental studies on the mechanisms of the placebo analgesic response, emotional activation in the treatment situation affects the magnitude of the placebo response (11).

The genetic effect on self-reported stress and systolic blood pressure (SBT) suggested that the affective correlates of pain may have different genetic predictors than reports of pain intensity. In the present data, 5-HTT, OPRM1 and COMT had main effects on reported stress regardless of whether the placebo treatment was provided. Furthermore, the A/A carriers in the placebo group, who reported the strongest placebo analgesic effect, also experienced a better stress relief than those with the */G variant of OPRM1. Interestingly, both 5-HTT and COMT had an interaction effect with group on the change in SBT from the pretest to the posttest, suggesting that the 5-HTT and COMT genotype could still influence the physiological response to pain.

Self-reported stress related to the affective component of pain might differ from objective measures of physiological activation (14, 41), and pain reports might differ from cerebral correlates of nociception (42). Thus, future studies investigating the impact of genetic variability could include more sophisticated physiological measures. Earlier data have suggested that decreased nociceptive activation induced by placebo treatment can be demonstrated by brain imaging (10), and that the OPRM1 genotype affects cerebral processes related to placebo responding (43). In the present study, however, we focused on the pain experience and demonstrate that genotyping of OPRM1 may be important to identify placebo responders. In conclusion, the OPRM1 A/A genotype is associated with increased placebo responses in healthy subjects free of clinical pain. We suggest that knowledge of genetic factors such as OPRM1 rs1799971 A/A may help improve designs for future RCTs in pain patients.

## Acknowledgments

The authors wish to thank Espen Bj0rkedal, Ph.D and Dagfinn Matre, Ph.D for providing the pain calibration procedure and the university hospital pharmacy at the University Hospital of Northern Norway for managing the production of the placebo and lidocaine/prilocaine creams used in this study. The present study was funded by The University of TromsØ, the Norwegian Institute of Occupational Health (STAMI), and The Arctic University of Norway, and a grant from the Northern Norway Health Authority to PerM. Aslaksen (grant number PFP1140-13).

## Materials and Methods

### Participants

The experiment included a total of 327 healthy Caucasian participants with a mean age of 23 years (SD = 4), 200 (61.2%) of whom were women. Participants were recruited by flyers on the campus of the University of Troms0, Norway, in the period 01.08.2013-01.08.2015. The study protocol was approved by the Regional Committee for Research Ethics in Health Sciences and Medicine, project number 2013/966. The participants signed an informed consent where they stated that they had no history of ongoing disease or any history of serious disease. Volunteers that used any type of prescribed medications or any type of analgesic medicine or therapy were not included in the study. Pregnant women were not allowed to participate. The participants were informed in the consent that the experiment tested the genetic influence of the effect of a commonly used local anesthetic cream. All participants received a gift card worth 200 NOK (approx. 25 USD) for reimbursement of expenses due to their participation in the study.

### Study Design

The design of the study was an experimental design with repeated measurements, consisting of a calibration procedure, pretest and posttest. Participants were randomized into three groups: The placebo group that got a moisturizing cream with no analgesic properties (E-45, Crookes Healthcare, UK), the natural history group receiving no treatment during the procedure, or the lidocain-prilocain cream group that received a commonly used local anesthetic cream (Emla, AstraZeneca, Norway). The group receiving the Emla cream was employed in the design to assure blinding of the experimenters, and these data were not used in the final analyses. Thirty-one (9.4%) ofthe participants were randomized into the lidocain-prilocain cream group. Thus, a total of 296 participants were used for the final analyses. The sample size was estimated by findings in a previous study performed on a Norwegian population (28), where approx. 25% were OPRM1 */G carriers, whereas 75% were A/A carriers. In order to obtain group sizes to include an adequate number of */G carriers, > 250 participants had to be included. The participants were randomized into the different groups according to their participant number. The experiment was executed according to a double-blind procedure in the placebo and Emla conditions where application of a placebo or Emla was required. The university hospital pharmacy at the University Hospital of Northern Norway produced 100-mL tubes of Emla cream (AstraZeneca, London, United Kingdom) and placebo cream (E45 Cream; Crookes Healthcare, Nottingham, United Kingdom). All tubes were numbered according to a list of codes and had an identical design. The code list was created by the university hospital pharmacy and was kept by the supervisor of the study, who did not participate directly in the experimental work. We chose the E45 cream as the placebo cream based on its similarities to Emla in color, odor, and consistency. A dose of 3 g of Emla or placebo was used for each participant, similar to Aslaksen et al (44).

The experiment occurred inside a steel cubicle (2.8 X 2.8 m) where the participants were placed in a comfortable chair. The cubicle was shielded from sound and electricity, and the temperature was kept at 20 °C. We applied thermal stimuli to the left underarm to induce pain. To assure an equal pain level across participants at the start of the experiment, a calibration procedure was performed. The calibration procedure estimated the stimulation intensity in °C sufficient to evoke a pain intensity of 60 on a 100-point computerized visual analog scale (VAS). In order to approximate the stimulus intensity needed to produce a rating of 60 on the VAS, we predicted the stimulus intensity by using Stevens’s power equation (45) VAS=b(t-t_0_)^c^. In this equation b is a scaling factor, t is the stimulus temperature, to is the intercept where VAS is assumed to be zero which was set to 35°C, and c is the exponent which defines the shape of the stimulus response function which was estimated based on the 10 calibration trials (46). The individually calibrated temperature was used throughout the experiment for each participant.

After calibration, the participants received two pain stimulations in the pretest. The duration of the stimulations (pretests and posttests) were 10 seconds from when the thermode reached the calibrated target temperature (43°C-47°C) until the start of the return to baseline at 32°C. The temperature of the thermode increased/decreased by 10°C/second. The interval between pre-test 1 and pre-test 2 was 30 seconds. The post-tests 1, 2 and 3 had the same temperature, duration and intervals as the pre-tests. Immediately after the pretest, the information about the treatment was provided to the participants allocated to the treatment groups where they received either placebo or Emla. The participants in the placebo group were told, “the cream that will be applied to your arm reduces pain. The substance in the cream is used as a local anesthetic in many pain-reducing remedies and is effective in the treatment of heat pain”. The participants were also told that there would be a break for a few minutes to allow the cream to produce the analgesic effect. In the natural history group, no cream was applied, and no information about the treatment was provided. The participants were told that there would be a break of a few minutes and that they could relax and wait. Systolic blood pressure and measures of perceived stress were obtained because reduction in these measures are shown to be associated with successful induction of placebo analgesia (11). Blood pressure was measured before the calibration procedure, after the treatment information was provided, and after the last posttest. Subjective stress was measured on a numerical rating scale with a range from 0 to 100 before the calibration procedure, after the pretests, after the treatment, and after the last posttest. The stress measurement was performed similar to previous studies (44, 47). The group of experimenters consisted of four females and two males with a mean age 24 years. The experimenters were psychology students who had extensive experience in performing experimental lab-procedures on human subjects. Three experimenters performed each experimental run, thus each participant interacted with three experimenters. The experimental procedure had a total duration of approximately 45 min.

### Genetic analyses

Collection of saliva and extraction of genomic DNA was done using an Oragene RNA sample collection kit (DNA Genotech Inc. Kanata, Ontario, Canada) according to the manufacturer’s instructions. SNP genotyping was carried out using predesigned TaqMan SNP genotyping assays for OPRM1, 5-HTT and COMT (Applied Biosystems, Foster City, CA, USA).

Approximately 10 ng genomic DNA was amplified in a 5-μl reaction mixture in a 384-well plate containing 1x TaqMan genotyping master mix (Applied Biosystems) and 1x assay mix, the latter containing the respective primers and probes. The probes were labeled with the reporter dye FAM or VIC to distinguish between the two alleles. After initial denaturation and enzyme activation at 95 °C for 10 min, the reaction mixture was subjected to 60 cycles of 95 °C for 15 sand 60 °C for 1 min. The reactions were performed on an ABI 7900HT sequence detection system. Negative controls containing water instead of DNA were included in every run. Genotypes were determined using the SDS 2.2 software (Applied Biosystems). Approximately 10% of the samples were re-genotyped, and the concordance rate was 100%.

To determine the length of the polymorphic promoter region of the 5-HTT, the DNA sequence was first amplified by polymerase chain reaction (PCR) and then separated by gel electrophoresis. PCR was carried out in a total volume of 25 μl containing ~60 ng of genomic template, 6.25 pmol of each primer and 1x Taq DNA Polymerase Master Mix (VWR international, Dublin, Ireland). The forward primer sequence was 5’ -GGCGT TGCCG CTCTG AATGC- 3’ and the reverse primer sequence was 5’ -GAGGG ACTGA GCTGG ACAAC CAC- 3’ (DNA technology A/S, Risskov, Denmark). Samples were amplified on a Perkin Elmer GeneAmp PCR 2400 system following an initial denaturing step for 3 min at 95 °C. The amplification consisted of 40 cycles including denaturing at 95 °C for 40 s, annealing at 60 °C for 20 s and elongation at 72 °C for 80 s, as previously described. The described PCR yielded a long (529 bp) and a shorter (486 bp) fragment (17). After four hours of separation at 100 Von a 2.5% agarose gel (MetaPhor Agarose, Lonza cologne GmbH, Cologne, Germany), Ge!Red dye was added, and the fragments were visualized by UV light (Biotium Inc, California, USA). A PCR 100-bp low ladder (Sigma-Aldrich CO, St. Louis, Mo, USA) was used to determine the length of the fragments.

### Statistical analyses

Continuous data were analyzed with linear mixed models (LMM) in SPSS version 23 (IBM, SPSS, USA). Group (placebo, natural history), sex of the participant, OPRM1, 5-HTT, COMT, and Trial were used as fixed factors. LMM was chosen because this method is suitable for analyzing data with unequal group sizes, handles missing data without losing power in the analyses compared to standard general linear models, and allows combinations of both fixed and random effects. The participants were assumed to exhibit significant individual variance, and the individual variance was treated as the only random effect in the repeated measures analysis. The p-values for comparisons within interactions were adjusted for multiple comparisons with Bonferroni corrections. To analyze the predictive value of OPRM1 on placebo responding, a logistic regression analysis was conducted. An alpha value of .05 was used in all analyses.

